# Enlarging viral mutation estimation: a view from the distribution of mutation rates

**DOI:** 10.1101/2025.03.20.644362

**Authors:** Taimá Naomi Furuyama, Isabel Maria Vicente Guedes de Carvalho, Luiz Mário Ramos Janini, Fernando Antoneli

## Abstract

The problem of empirical estimation of mutation rates is fundamental for the understanding of viral evolution. The estimation of viral mutation rates is based on varied and often complex methods carried out through experiments essentially designed to count mutation frequencies. Mutation rates are defined as the probabilities of nucleotide substitutions, typically reported as a single number in units of mutation (substitution) per base (nucleotide) per replication cycle or per cell infection, depending on the replication mode of the virus. Even more, the uncertainty quantification of these estimates is so difficult that it is rare to find it reported in the literature. The values for the same virus reported in literature fall within a broad range, sometimes spanning two orders of magnitude. For instance, the mutation rates range from 10^*−*8^ to 10^*−*6^ mutation per base per cell infection for DNA viruses and from 10^*−*6^ to 10^*−*4^ mutation per base per cell infection for RNA viruses. In this paper, we propose an alternative perspective on the estimation of mutational rates, which avoids the use of consensus sequences and/or serial passages. Our approach leverages the large amount of sequencing data produced by high throughput sequencing technologies coupled to an experimental design that performs a single replication cycle from an initial clonal viral population. We propose to replace the single numeric mutation rate with a distribution of mutation rates (DMR), together with a procedure to implement the estimation of this distribution from sequencing data and show that it can be estimated from sequencing data. Even though the focus of this paper is the development of the approach centered on the DMR it is straightforward to produce point and interval estimates of the mutation rates, including uncertainty quantification. In addition to the estimation of the DMR, we provide a theoretical characterization of it, as being well-approximated by a log-normal distribution. Finally, we study some non-trivial properties of the DMR related to a remarkable invariance under down-scaling the distribution from the genome to its subunits.

## 1 Introduction

A key concept in viral population knowledge is mutation rates. It is necessary to understand the processes where a drug can act and its risks/side effects, like the pro-drug molnupiravir that leads SARS-CoV-2 to lethal mutagenesis through nucleoside analogs or hepatitis C virus (HCV) NS5A and NS5B inhibitors [8, 20, 55]. Moreover, Influenza virus strain recommendations for seasonal vaccination worldwide and COVID-19 variant classification to identify the impact risk on the epidemiological situation are public health decision tools based on the dynamics of viral populations, including their mutations [9, 18].

Spontaneous mutations are mutations that occur in the absence of exogenous agents. They may be due to errors made by DNA polymerases during replication or repair, errors made during recombination, the movement of genetic elements, or spontaneously occurring DNA damage. The rate at which spontaneous mutations occur is a fundamental evolutionary parameter.

The mutation rate of a virus is the expected number of mutations that a viral particle will sustain during its replication cycle. In empirical experiments designed to estimate spontaneous mutation rates, it is very common to measure mutation frequencies which, together with other information, is used to compute a single number *µ*, the *mutation rate estimate* [10]. Usually, *µ* is obtained as the mean or the median of some sampling-counting scheme, and in rare situations, it comes along with an uncertainty measure, e.g., the standard deviation or confidence interval [11].

Indeed, accurate and precise estimates have proved challenging, because of the difficulty in sampling genomic data and counting mutation frequencies and the many possibilities of how to combine mutation frequencies and population history to calculate mutation rates. Because of these difficulties, the literature is fraught with apparently disparate estimates – typically spanning one or two orders of magnitude. For instance, it is possible to find in the literature the following estimates for three important viruses: (i) HIV, 5.4 × 10^−5^ m/b/r (mutations/base/replication cycle) and 3.0 × 10^−4^ m/b/r (purified HIV-1 RT) [12], 3.4 × 1 0^−5^ m/b/r [30], 2.4 × 1 0^−5^ m/b/c (mutations/base/cell infection) [42], 1.4 × 1 0^−5^ m/b/r and 5.9 × 1 0^−4^ m/b/r (purified HIV-1 RT) [1], 6.2 × 1 0^−5^ m/b/c [7]; (ii) HCV, 1.2 × 1 0^−4^, 2.5 × 1 0^−5^, 2.0 × 1 0^−5^, 3.5 × 1 0^−5^ m/b/c; (iii) Influenza virus A, 7.1 × 1 0^−6^, 4.5 × 1 0^−5^, 3.9 × 1 0^−5^, 3.1 × 1 0^−5^ m/b/c [41].

Mutational rates are composed of diversified processes according to the virus. It is relevant to consider the enzyme fidelity, whether the polymerase or the exonuclease, the replication process, and its interaction with the immune response system of the host, nucleic acid spontaneous damage, access to post-replicative repair, number of available nucleotides, quality, and structure of emerging templates [41].

In this paper, we propose an alternative perspective on the estimation of mutational rates, which avoids the use of consensus sequences and/or serial passages. Our approach leverages the large amount of sequencing data produced by high-throughput sequencing technologies coupled with an experimental design that performs a single replication cycle from an initial clonal viral population [33]. With this, we expect to bypass the difficulties with sampling and/or counting mutation frequencies and account for all mutation occurrences, including possible *de novo*, neutral, deleterious, and lethal mutations [34]. Hence, to account for the wide range of mutation rates in a single genome, we propose to model them as a ‘distribution of mutation rates (DMR)’ rather than as a single numerical value.

We should mention that other authors already considered the notion of distribution of mutation rates in different contexts: [2] for viral populations and [50] for microbial populations. However, to the best of our knowledge, we are the first to propose a method for the distribution of mutation rates estimation based on high throughput sequencing data.

Mutational rates defined as a single numerical value can hide potentially useful information. A viral population is dynamic and uses genomic strategies to reach its balance between mutations, drift, and selection according to the fitness landscape. The main novelty of this paper is the proposal to replace the single numeric mutation rate for a distribution of mutation rates (DMR), with a procedure to implement the estimation of this distribution from sequencing data. This ‘population’ viewpoint allows for a more comprehensive framework for the representation of the mutational dynamics, possibly associated with the dynamics of a virus quasispecies.

## 2 Materials and Methods

### 2.1 Theoretical background: the distribution of mutation rates

The motivation to introduce the distribution of mutation rates (DMR) is to reflect the fluctuations of mutation rates across co-existing individuals. Therefore, one is led to consider the mutation rate per locus as a random variable.

Because of this shift in the perspective, this approach allows one to see the effects of population heterogeneity in the mutation rate, that is, variation in the population mean mutation rate. This approach contrasts with many previous studies that compared the presence versus absence of a ‘mutator’ type, thus changing the mean and the variance of mutation rate [2].

We propose to model the *distribution of mutation rates (DMR)*, in the context of RNA viral populations, as a log-normal probability distribution to account for the following properties:

1. most mutation rates are low and very close to 0, around 10^−4^ s/n/c,
2. none of them are zero and almost none are extremely low, e.g. ≤ 10^−6^ s/n/c,
3. very few of them are higher.

Since a log-normal distribution has a long tail of few high-rate mutations it is expected that a DMR has Mean *>* Median *>* Mode. This means that using the DMR to obtain point estimates of the mutation rate as mean or median values can give higher values than traditionally obtained, as the usual methods for estimating mutation rates are sensitive to extreme cases encountered in mutational hotspots.

### 2.2 Estimation of the distribution of mutation rates

The sequencing data used in this paper was generated in a set of single replication cycle experiments with HIV-1. We needed only the publicly available data published in [33]. In a very summarized way, the cell culture procedure was performed through transfection by two types of pseudoviruses (CCR5 or CXCR4), two types of cells (T cells or PBMCs), and two cellular states (stimulated and non-stimulated). The viral sequences were obtained from the proviral genome using the SOLiD technology. The assembly was obtained by mapping the reads against a reference and the whole genome (without the LTR regions) is given by positions 791 to 9,085. The plasmid DNA used to construct the pseudoviruses was sequenced, as well, as generating the control (CTRL) experiment. Totalizing 7 experimental conditions and a control experiment: (1) NS_CD4_R5, (2) NS_CD4_X4, (3) NS_PBMC_R5, (4) NS_PBMC_X4, (5) S_CD4_R5, (6) S_PBMC_R5, (7) S_PBMC_X4, (8) CTRL (a putative experimental condition ‘S_CD4_X4’ is missing because the sequencing was not successful in this case).

For the data analysis, an *Empirical Bayes* approach was used, with a control state (the pseudovirus genome) as the prior condition and the obtained sequences as an *a posteriori* condition [32]. This analysis generated estimates for the multinomial probability distribution of 4 nucleotides at each site: p_A, p_T, p_G, and p_C, with p_A+p_T+p_G+p_C= 1 [56].

From the monomial probability distributions, one can compute the *complementary probability* at each site, defined as

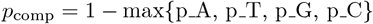

The interpretation of *p*_comp_ is that it is the sum of the probabilities of the 3 remaining nucleotides after the most probable is removed. The complementary probability *p*_comp_ is an overall indicator of the fluctuation of the possible nucleotide at each site. A higher *p*_comp_ indicates that two or more nucleotides are highly likely at that position. A lower *p*_comp_ indicates that a single nucleotide is overwhelmingly prevalent at that position.

The distribution of mutation rates (DMR) was estimated as the empirical distribution generated by the set of the complementary probabilities {*p*_comp_} for all sites. Since we expected that the empirical DMR approximates a log-normal distribution it was natural to transform the original data {*p*_comp_} to a logarithmic − scale, which should follow a normal or Gaussian distribution. We use the logarithm transformation *s* = − log(*p*) also known as the *s* score or *surprisal*, or *self-information* [6, 16, 17, 43, 45]. *In this paper, we will always use the natural logarithm, denoted by ‘log’ and its corresponding exponential function denoted by ‘exp’*.

Finally, we applied an exclusion criterion to disregard sites that (i) were extremely variable and (ii) where the uncertainty of the estimate of max{p_A, p_T, p_G, p_C} were below the accuracy of the method (see [56]) – this means that, for these sites, *p*_comp_ is so small that we cannot distinguish it from 0. The first case is obtained by excluding sites whose complementary probability was below 6.0 × 1 0^−3^ and the second case is obtained by excluding sites whose complementary probability was above 10^−4.2^. Despite this exclusion of sites, about 94% of the initial sites were retained, resulting in approximately 7,800 sites per genome.

The estimated DMR and log-transformed DMR (log-DMR) are univariate empirical distributions and hence can be visualized in a simple 2D plot. Moreover, several measures of central tendency and dispersion, such as mean, median, mode, range, upper and lower quartiles, variance, and standard deviation, can be computed for these distributions. In particular, the mean, median, and mode can serve as point estimates for mutation rate seen as a single numerical value, for the sake of comparison with the estimates from the literature.

All the computations were carried out using the statistical software R with the Rstudio IDE [36, 37]. The R package ModeEst (Mode Estimation, version 2.4.0 [35]) was used to perform the mode estimation. The R package poweRlaw (Analysis of Heavy Tailed Distributions, version 0.80 [13, 14]) was used to perform the log-normal fit and bootstrap procedure for the hypothesis test using the PLFit algorithm [5]. The R package Mass (Modern Applied Statistics with S, version 7.3-54, [38]) was used to compute the Q-Q plots and to obtain the maximum likelihood fit for the univariate normal distributions.

### 2.3 Characterization and properties of the distribution of mutation rates

We performed a pipeline of exploratory data analysis on the empirical DMRs obtained from the 7 experimental conditions and the control. The pipeline consists of 2 main steps: (a) pairwise two-sample Kolmogorov-Smirnov tests comparing the DMRs from the 7 experimental conditions and the control to asses the similarity or dissimilarity between them, and (b) a robustness analysis that comprised excluding certain data to verify whether any clear pattern or genomic hotspots could be identified. This pipeline was applied to the whole genome, specific genes, and fragments.

#### 2.3.1 Pairwise two-sample Kolmogorov-Smirnov test

We performed a pairwise two-sample Kolmogorov-Smirnov test with all experimental conditions and the control. The goal was to verify whether the empirical DMRs were the same or not. The test was performed for all possible pairs of 8 conditions resulting in a total of *K* = (8 × 7)*/*2 = 28 tests. We assumed a significance level of α = 0.05 per test, hence, using the *Bonferroni adjustment*, we had a total significance level of α_B_ = 0.001.

#### 2.3.2 Robustness analysis

Recall that we have excluded sites that (i) were extremely variable and (ii) had uncertainty below the accuracy of the estimation method. It is worth mentioning that the hyper-mutated sites, excluded by criterion (i), were already analyzed by [33]. The sites that satisfy exclusion criteria (ii) can be considered as the ‘invariant sites’, since for them *p*_comp_ is statistically indistinguishable from 0.

Therefore, we investigated if there were some patterns of invariant sites that were recurrent among the experimental conditions. We verified if the invariant sites were concentrated or randomly spread across the genome. We computed the sets of invariant sites of each experimental condition and determined the invariant sites that were excluded from more than one experimental condition. For those invariant sites that appeared more than once we counted in how many conditions it was present. Using this information we also searched for possible mutational hotspots.

Finally, to verify whether a significant number of consecutive sites or long-length fragments were excluded when applying the exclusion criteria, we computed the length and position of the gaps left after the exclusion of sites.

## 3 Results

### 3.1 Distributions of mutation rates

We estimated the empirical DMRs for all 7 experimental conditions and the control. We also computed a ‘consensus’ empirical DMR by averaging the 7 experimental conditions over each site. In the following we will discuss the consensus and the control in more detail, showing that the individual experimental conditions are very similar (see the Supplementary Material).

In Figures 1 and 2 we show 4 plots obtained from the consensus empirical DMR and the control empirical DMR. In the top left panels, we show the histograms of the empirical DMRs. In the bottom left panels, we show the histograms of the corresponding log-DMRs. In the bottom right panels, we show the Q-Q plot of the log-DMRs. In the top right panels, we show the corresponding empirical cumulative distribution in log-log scale with a log-normal tail fit [14].

**Figure 1:**
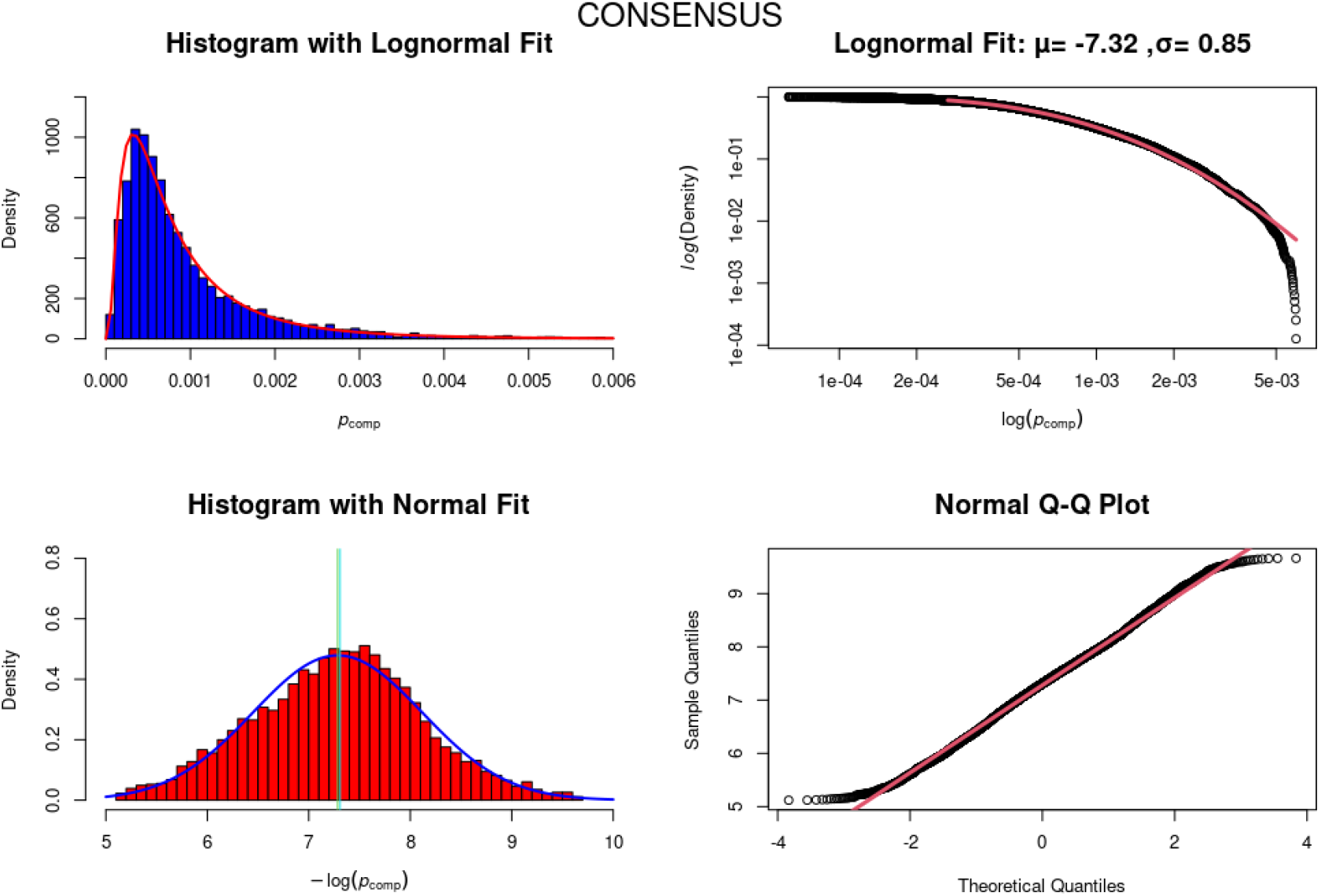
Different views of the consensus empirical DMR. (Top Left) Histogram of the empirical DMR, with the theoretical log-normal distribution (red curve) superimposed. (Bottom Left) Histogram of the empirical log-DMR, with the theoretical normal distribution (blue curve) superimposed. The green and cyan vertical lines indicate the mean and median, respectively. (Top Right) Empirical cumulative DMR in log-log scale with log-normal tail fit (red curve). (Bottom Right) Q-Q plot of the empirical log-DMR. Here, the red curve indicates the expected graph if the empirical distribution is exactly the same as the theoretical.

**Figure 2:**
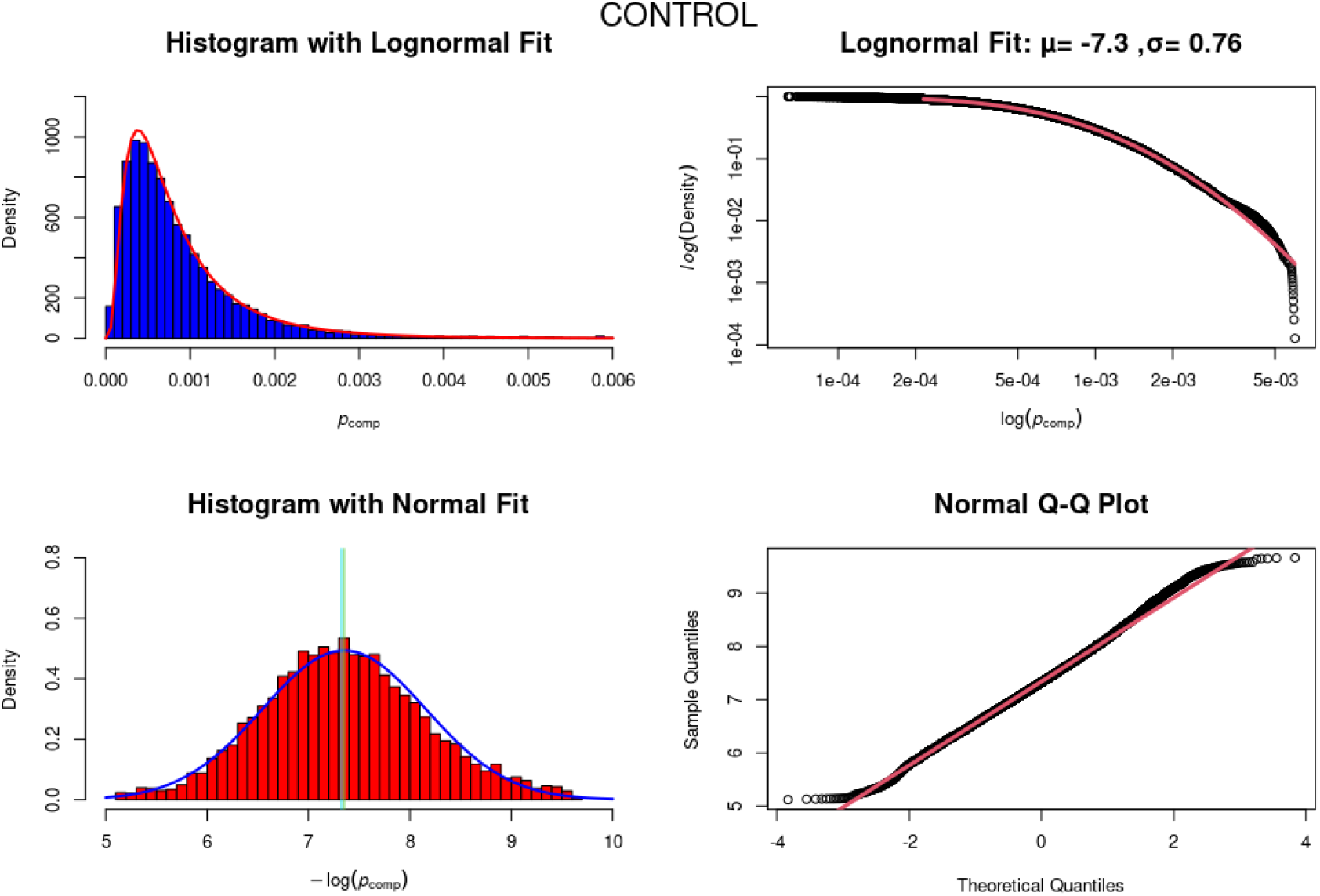
Different views of the control empirical DMR. (Top Left) Histogram of the empirical DMR, with the theoretical log-normal distribution (red curve) superimposed. (Bottom Left) Histogram of the empirical log-DMR, with the theoretical normal distribution (blue curve) superimposed. The green and cyan vertical lines indicate the mean and median, respectively. (Top Right) Empirical cumulative DMR in log-log scale with log-normal tail fit (red curve). (Bottom Right) Q-Q plot of the empirical log-DMR. Here, the red curve indicates the expected graph if the empirical distribution were exactly the same as the theoretical.

All these plots are meant to show that the empirical DMRs indeed follow a log-normal distribution. The histograms on the left are superimposed by the hypothesized theoretical distribution. The Q-Q plot and the log-normal tail fit show that the empirical DMR is very well approximated by a theoretical log-normal distribution. In addition, we performed the bootstrap procedure of [14] to test the null hypothesis (H_0_) that the data is generated from a log-normal distribution against the alternative hypothesis (H_1_) that the data is not generated from a log-normal distribution. In all cases, we obtained *p >* 0.9 indicating that the null hypothesis cannot be rejected.

It is important to stress that based on any given finite sample, there is absolutely no way to tell how likely the hypothesis is that it was sampled from a log-normal, or any heavy-tailed distribution, in fact [48]. The best thing one can hope to achieve is that a log-normal can acceptably explain the data – as we did above using the bootstrap procedure of [14].

It is interesting to note that the empirical DMR of the control condition also seems to be well approximated by log-normal. We will add more comments to this observation as we proceed further in our analysis.

Table 1 shows some summary statistics of the DMRs for all 7 experimental conditions, the control, and the consensus. As expected from a log-normal distribution, we always have that Mode *<* Median *<* Mean. Moreover, the interval [Mode, Median] is a robust interval estimate for the mutation rate. Here, we consider the interval between the Mode and the Median because these two central tendency measures are more robust than the Mean, which is very sensitive to variation in the tail of the distribution. Given the large variability of the point estimates of the mutation rate from the literature, perhaps a more inclusive possibility is to use the interval associated with the interquartile range (IQR), defined as [Q_1_ – Q_3_], as a robust interval estimate for the mutation rate.

**Table 1:** Summary statistics of the empirical DMR for the 7 experimental condition, the control, and the consensus. Min.: minimum value; Q_1_: first quartile; Q_3_: third quartile; Max.: maximum value; R.S.: proportion of retained sites, i.e. the sites that were considered for the analysis.

| Condition | Min. | Q<sub>1</sub> | Mode | Median | Mean | Q<sub>3</sub> | Max. | R.S. |
| --- | --- | --- | --- | --- | --- | --- | --- | --- |
| CONTROL | $6.38 \times 10^{-5}$ | $3.82 \times 10^{-4}$ | $4.15 \times 10^{-4}$ | $6.59 \times 10^{-4}$ | $8.79 \times 10^{-4}$ | $1.10 \times 10^{-3}$ | $5.98 \times 10^{-3}$ | 95% |
| NS_CD4_R5 | $6.38 \times 10^{-5}$ | $3.85 \times 10^{-4}$ | $3.93 \times 10^{-4}$ | $6.61 \times 10^{-4}$ | $9.39 \times 10^{-4}$ | $1.17 \times 10^{-3}$ | $5.97 \times 10^{-3}$ | 95% |
| NS_CD4_X4 | $6.33 \times 10^{-5}$ | $3.74 \times 10^{-4}$ | $3.99 \times 10^{-4}$ | $6.51 \times 10^{-4}$ | $9.40 \times 10^{-4}$ | $1.18 \times 10^{-3}$ | $5.87 \times 10^{-3}$ | 93% |
| NS_PBMC_R5 | $6.34 \times 10^{-5}$ | $3.94 \times 10^{-4}$ | $4.37 \times 10^{-4}$ | $6.64 \times 10^{-4}$ | $9.50 \times 10^{-4}$ | $1.17 \times 10^{-3}$ | $5.99 \times 10^{-3}$ | 95% |
| NS_PBMC_X4 | $6.35 \times 10^{-5}$ | $3.83 \times 10^{-4}$ | $3.95 \times 10^{-4}$ | $6.45 \times 10^{-4}$ | $9.17 \times 10^{-4}$ | $1.13 \times 10^{-3}$ | $5.99 \times 10^{-3}$ | 95% |
| S_CD4_R5 | $6.32 \times 10^{-5}$ | $3.49 \times 10^{-4}$ | $3.22 \times 10^{-4}$ | $6.13 \times 10^{-4}$ | $8.80 \times 10^{-4}$ | $1.10 \times 10^{-3}$ | $5.98 \times 10^{-3}$ | 93% |
| S_PBMC_R5 | $6.31 \times 10^{-5}$ | $3.52 \times 10^{-4}$ | $3.98 \times 10^{-4}$ | $5.98 \times 10^{-4}$ | $8.89 \times 10^{-4}$ | $1.09 \times 10^{-3}$ | $5.95 \times 10^{-3}$ | 94% |
| S_PBMC_X4 | $6.33 \times 10^{-5}$ | $3.53 \times 10^{-4}$ | $4.03 \times 10^{-4}$ | $6.20 \times 10^{-4}$ | $9.36 \times 10^{-4}$ | $1.14 \times 10^{-3}$ | $5.97 \times 10^{-3}$ | 94% |
| CONSENSUS | $6.34 \times 10^{-5}$ | $3.70 \times 10^{-4}$ | $3.92 \times 10^{-4}$ | $6.36 \times 10^{-4}$ | $9.22 \times 10^{-4}$ | $1.14 \times 10^{-3}$ | $5.96 \times 10^{-3}$ | 94% |

Passing to the log-transformed DMRs (log-DMRs), Figure 3 shows the box plot of the log-DMRs for the 7 experimental conditions and the control. Since these distributions are expected to be Gaussian, we have that Mode Median Mean. In order to quantify how much divergence there is between the Mode, the Median, and the Mean we computed the skewness measure of each distribution. In a certain sense, this also quantifies how much the empirical distribution deviates from a theoretical Gaussian distribution.

**Figure 3:**
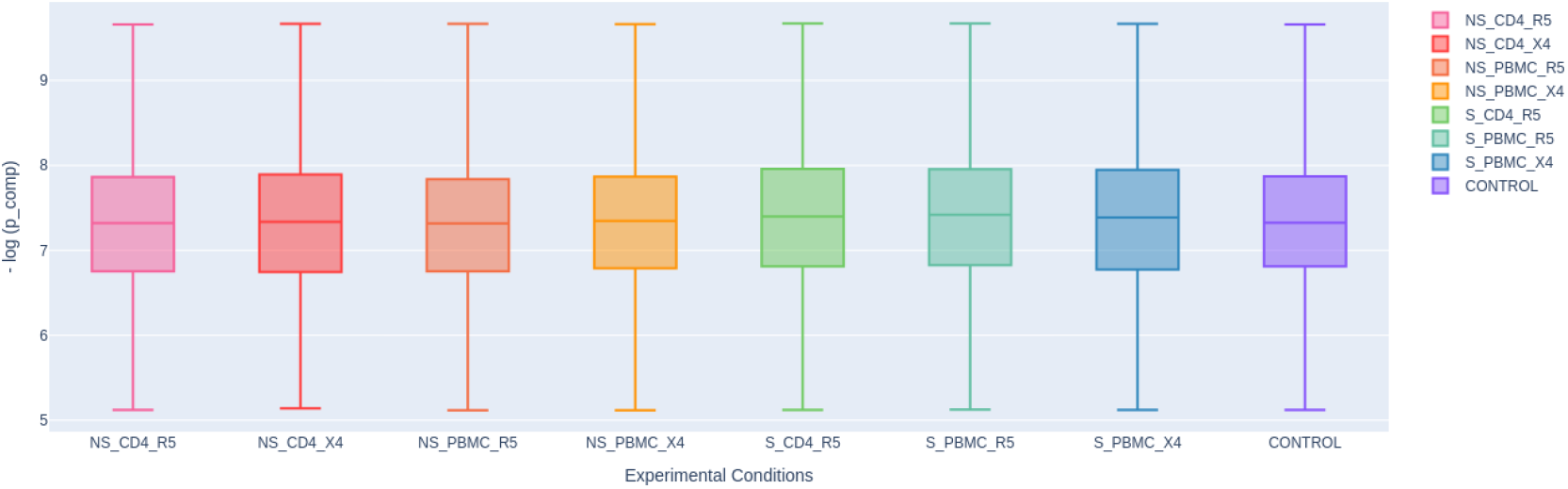
Box plots of the log-DMRs for the 7 experimental conditions and the control. The whiskers were set to be the minimum and the maximum.

The *skewness* of a distribution is a measure of the asymmetry of the distribution about its mean. If the distribution is both symmetric and unimodal, then Mode = Median = Mean. More generally, for unimodal distributions, the skewness value can be positive, zero, negative, or undefined. Roughly speaking, negative skewness indicates that the left tail is longer and the mass of the distribution is concentrated on the right side; positive skewness indicates that the right tail is longer and the mass of the distribution is concentrated on the left side.

We estimated the skewness of our data for every experimental condition using the functions for skewness estimation of the package ModeEst [35]. The result is shown in Table 2. Overall, the deviation from a perfectly symmetric distribution is very small in all cases except the control, which was the highest value of all. Typically, when the skewness is between ™0.5 and 0.5 the distribution is considered almost symmetric. Nevertheless, observe that the higher values are from the non-stimulated conditions and the control and the lower values (even negative) are from the stimulated conditions and the consensus.

**Table 2:** Skewness estimates for the 7 experimental conditions, the control and the consensus. The stimulated (S) conditions and the consensus had lower or negative values whereas the non-stimulated (NS) ones and the control had higher.

| Condition | Skewness | Condition | Skewness |
| --- | --- | --- | --- |
| NS_PBMC_X4 | 0.046 | S_PBMC_X4 | -0.064 |
| NS_PBMC_R5 | 0.031 | S_PBMC_R5 | -0.003 |
| NS_CD4_R5 | 0.043 | S_CD4_R5 | 0.016 |
| NS_CD4_X4 | 0.061 | - | - |
| CONTROL | 0.138 | CONSENSUS | 0.008 |

### 3.2 Exploratory Data Analysis

#### 3.2.1 Pairwise two-sample Kolmogorov-Smirnov test

For the pairwise two-sample Kolmogorov-Smirnov test, the null hypothesis is that both empirical distributions were sampled from the same population. The results are shown in Table 3. The most important observation is that the conditions are split into three groups: (i) the control, which is distinct from all other conditions, (ii) the stimulated conditions and (iii) the non-stimulated conditions. Two pairwise comparisons were marginally close to the rejection threshold: (1) CTRL × NS_PBMC_X4 and (2) S_CD4_R5 ×S_PBMC_X4.

**Table 3:** Pairwise two-sample Kolmogorov-Smirnov test with correction for multiple testing (number of tests *K* = 28). The significance level per test was set to *α* = 0.05 and using *Bonferroni adjustment* the total significance level was α_B_ = 0.001. The table shows the *p*-values of all pairwise tests. Two pairs were marginally close to the significance level, namely CTRL × C4 and C5 × C7. The conditions are C1: NS_CD4_R5, C2: NS_CD4_X4, C3: NS_PBMC_R5, C4: NS_PBMC_X4, C5: S_CD4_R5, C6: S_PBMC_R5, C7: S_PBMC_X4, CTRL: Control.

|  | CTRL | C1 | C2 | C3 | C4 | C5 | C6 | C7 |
| --- | --- | --- | --- | --- | --- | --- | --- | --- |
| CTRL | - | $5 \times 10^{-4}$ | $5 \times 10^{-5}$ | $3 \times 10^{-5}$ | $5 \times 10^{-3}$ | $1 \times 10^{-5}$ | $3 \times 10^{-8}$ | $3 \times 10^{-6}$ |
| C1 | | - | 0.2 | 0.6 | 0.07 | $5 \times 10^{-6}$ | $1 \times 10^{-8}$ | $2 \times 10^{-5}$ |
| C2 | | | - | 0.03 | 0.02 | $2 \times 10^{-4}$ | $2 \times 10^{-6}$ | $1 \times 10^{-3}$ |
| C3 | | | | - | 0.1 | $1 \times 10^{-8}$ | $1 \times 10^{-9}$ | $1 \times 10^{-7}$ |
| C4 | | | | | - | $6 \times 10^{-5}$ | $2 \times 10^{-6}$ | $5 \times 10^{-5}$ |
| C5 | | | | | | - | 0.5 | $5 \times 10^{-3}$ |
| C6 |  |  |  |  |  |  | - | 0.1 |
| C7 |  |  |  |  |  |  |  | - |

#### 3.2.2 Robustness analysis

We searched for patterns in the excluded dataset to reduce the possibilities for bias in our analysis. About 94% of all sites were considered for analysis, and 6% was discarded. The excluded sites were distributed throughout the genome, not indicating any preferential region of concentration. The positions of the gaps between the sites display an almost uniform distribution throughout the genome (see Figure 4(Right) for the consensus condition and **??** of the Supplementary Material for all other conditions). The distribution of gap lengths, shown in Figure 4 (Left) for the consensus condition, gives the proportion of gaps of length *ℓ* = 0, 1, …, *ℓ*_max_. We performed chi-square tests, for all conditions, showing that it follows a negative binomial distribution, confirming the randomness of gap clustering [47].

**Figure 4:**
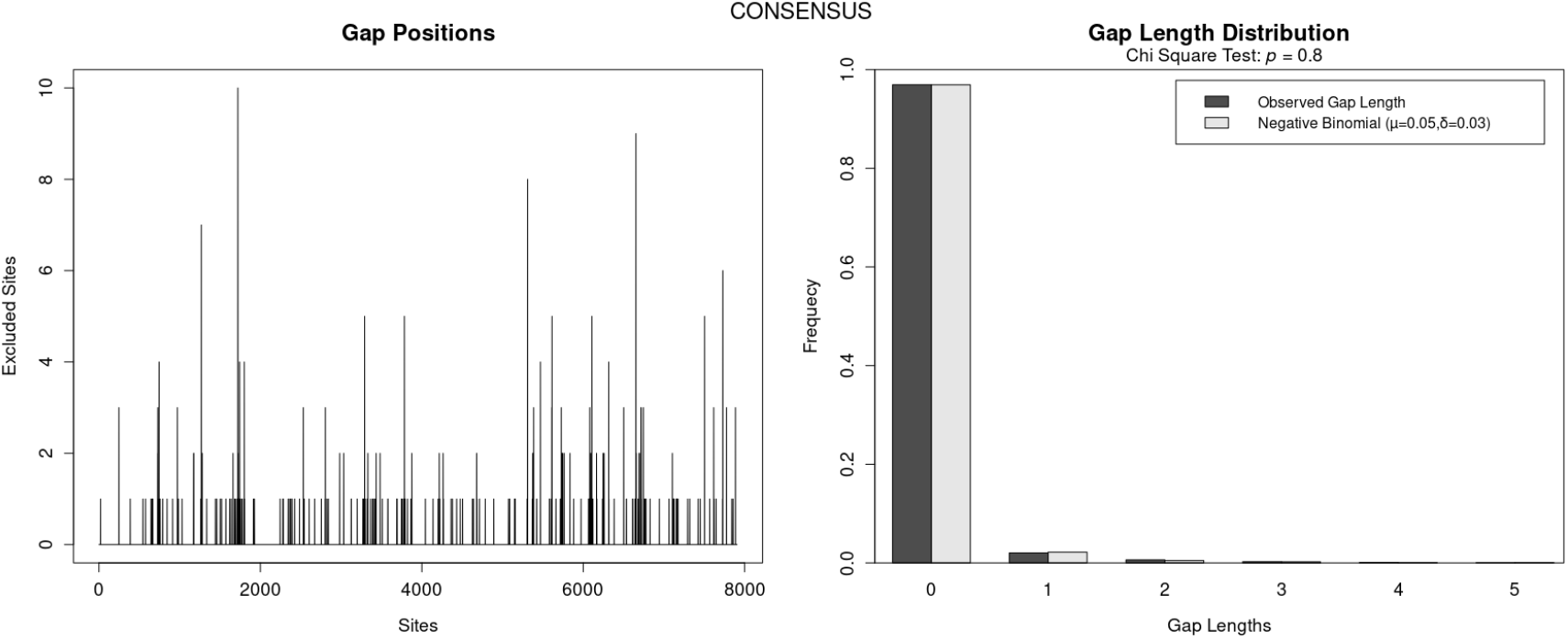
Gap positions and gap length distribution for the consensus. (Left) Gap positions on the genome. The length of the bar is the size of the gap. The marks in the *x*-axis represent the absolute number of sites in the specific sequence and not their location in the genome. (Right) Distribution of gap lengths. The panel shows the observed gap lengths (dark gray) and the expected negative binomial distribution (light gray) with parameters *µ* = 0.05 (mean) and *δ* = 0.03 (dispersion). For each length *ℓ* = 0, 1, …, 10 the bars give the proportion of sites with length *ℓ*. Here, the *x*-axis shows the lengths up to *ℓ* = 5.

Finally, we also verified if there were invariant sites that were recurrent in more than one condition. Considering the universe of analyzed sites and conditions, we found that less than 1% of the sites appeared in two or more conditions.

### 3.3 Scale invariance and information dimension

#### 3.3.1 Down-scaling of the distribution of mutation rates

The idea here is to investigate how the empirical DMR behaves under down-scaling from the genome to its subunits (e.g. genes, fragments, etc.).

Table 4 shows the summary statistics of the consensus empirical DMRs for the whole genome, genes, and smaller fragments, up to a length of 50 sites. The same result holds for all 7 conditions and the control (data not shown). Observe that there is a maintenance in the order of magnitude of the Mode, the Median, and the Mean, regardless of the length of the fragment, up to fragment length around (or below) 200 nucleotides.

**Table 4:**
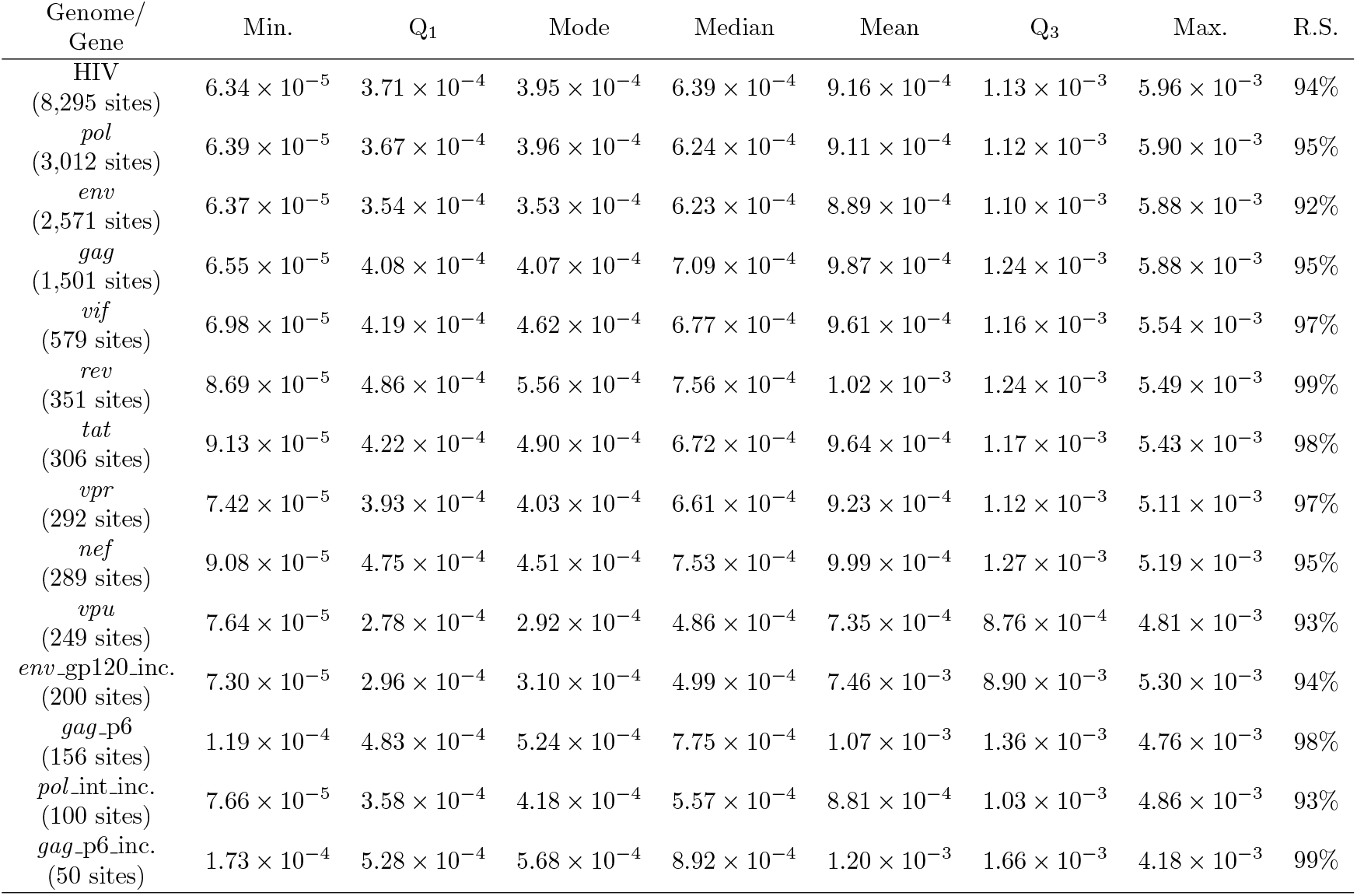
Summary statistics of the consensus empirical DMRs for the whole genome, genes, and smaller fragments, up to a length of 50 sites. Min.: minimum value; Q_1_: first quartile; Q_3_: third quartile; Max.: maximum value; R.S.: proportion of retained sites, i.e. the sites that were considered for the analysis.

This result suggests that the DMR is *scale-invariant* in the sense that the information contained in the distribution is (up to a certain limit) independent of the length of the genomic fragment that carries the information.

#### 3.3.2 Estimation of Information Dimension

To test if the DMR has some sort of invariance under down-scaling we resorted to the theory of information, more precisely, the notion of *information dimension*.

The idea of *information dimension* is to afford a measure of the ‘fractal character’ of a probability distribution associated with a given system. It characterizes the growth rate of the Shannon entropy given by successively finer discretizations, called coarse-grained approximations, of the state space of the system [39]. We provide a brief of the main concepts related to information theory and information dimension in the Supplementary Material, appendix **??**.

To apply the construction of information dimension to our context we start by partitioning the genome into *N* fragments of equal length, where *N* = 2, 3, 4, …, *N*_max_. Denote obtained the set of fragments, i.e. the partition of the genome into *N* parts, by 𝒜_*N*_ = {*A*_1_, …, 𝒜_*N*_}. Hence, if the length of the genome (i.e. the number of sites) is denoted by *L*, then the length of each fragment of A_*N*_ is approximately *L/N*.

Now we associate a discrete probability distribution *P*_*N*_ = *p*(*A*_*j*_) : *j* = 1, …, *N* to each partition 𝒜_*N*_ as follows. Let

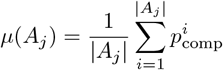

where |*A*_*j*_| is the number of sites in the fragment *A*_*j*_ and *p*^*i*^ is the complementary probability associated to the *i*-th site of the fragment *A*_*j*_. Then we define

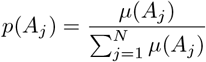

That is, *p*(*A*_*j*_) is the normalized average of the complementary probabilities of each fragment. Now we define the Shannon entropy of the partition 𝒜_*N*_ by

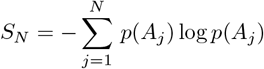

In this setting, we introduce the ‘scaling parameter’ *ε* = 1*/N* of the system and consider the sequence

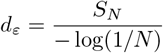

When *ε* decreases towards 0 it is expected that the sequence *d*_*ε*_ approximates the Shannon information dimension of the DMR. However, as we decrease *ε* (equivalently increases *N*) the fragments *A*_*j*_ become very small and the associated empirical DMR starts to degrade due to under-sampling. In fact, from Table 4 one can observe that when the length of the fragment is about 200, some of the summary statistics of empirical DMR start to diverge from the trends.

The intuition behind this definition is the following. If we interpret the sets *A*_*j*_ as a ‘box of size’ *ε* over the genome, then we can interpret *S*(*ε*) = *S*_*N*_ as the average information per box. Hence, the number *d*_*ε*_ is the exponent of the box size scaling of the average information needed to identify an occupied box, that is

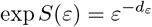

The exponential of the Shannon entropy exp *S* is called *perplexity* in the fields of machine learning and statistical modeling and is called *diversity index of order* 1 in the fields of ecology and demography [24].

In our case, *L* ≈ 7800 (i.e. the total number of sites after the application of exclusion criteria). Hence, if we set *N*_max_ = 20 the average fragment length is *L/N*_max_ = 390. Comparing this length with the results from Table 4 we expect to stay in the region where the trends of the summary statistics are preserved. However, if we set *N*_max_ = 100, thus *L/N*_max_ = 78 we expect to go through the region where the trends of the summary statistics start to diverge.

In Fig.5 we show the computation of the sequence *d*_*ε*_ as function of *ε* = 1*/N* for *N*_max_ = 10, 20, 50, 100. The data used in the computation was from the consensus condition. The other conditions give the same result. The estimate of the Shannon information dimension is 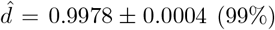. The fact that 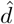 is very close to 1 corroborates the fact that the family of distributions 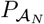 is very close to a continuous uniform distribution. This is strong evidence that the DMR is scale-invariant with scaling exponent ≈ 1.

We performed two tests to check the robustness of the previous result. The first test is accomplished by rearranging the sites in such a way as to break the natural sequential distribution of the complementary probabilities. This test shows that the localization of the mutation rates in the genome is crucial for the scale invariance to occur, in the sense that the ‘spatial’ distribution of *p*_comp_ along the genome is almost uniform. So much so that when we disrupted this ‘spatial’ distribution the scale invariance was lost. In Figure 6 we plot the same data as in Figure 5, except that we sorted the sites in increasing order concerning *p*_comp_. The regression line used to estimate the information dimension is no longer horizontal. The slope is non-zero with high significance (*p <* 10^−15^), and hence the estimate of *d* by the intercept of the regression line does not make sense anymore.

**Figure 5:**
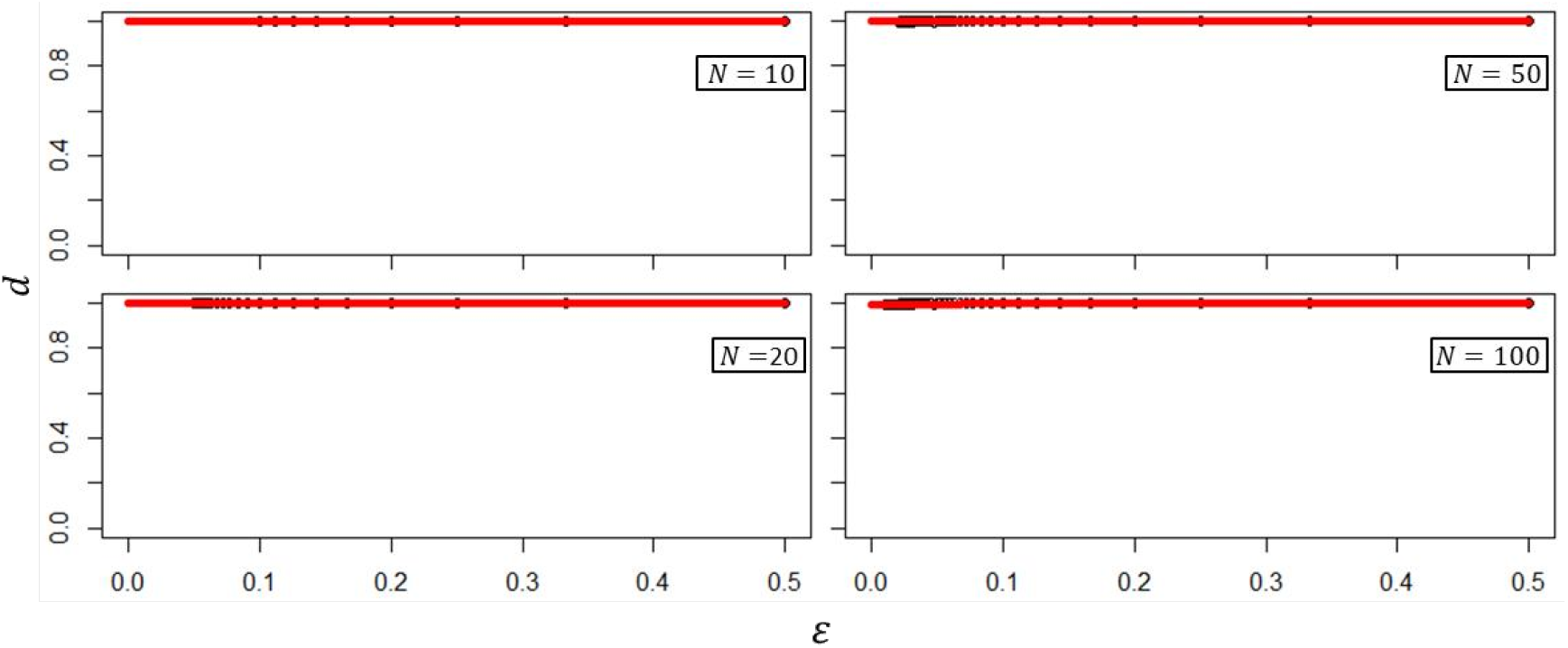
Shannon information dimension *d*_*ε*_ as function of the scaling parameter *ε* = 1*/N* for *N*_max_ = 10, 20, 50, 100. The data shown is obtained from the consensus condition. The other conditions give the same result. The red line is obtained by linear regression with a slope estimate of 0.003 (*p >* 0.001) and an intercept estimate of 0.9978 (*p <* 10^−15^) Hence, the regression line is parallel to the horizontal axis. The estimate of the information dimension is given by the intercept estimate 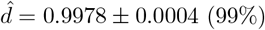.

**Figure 6:**
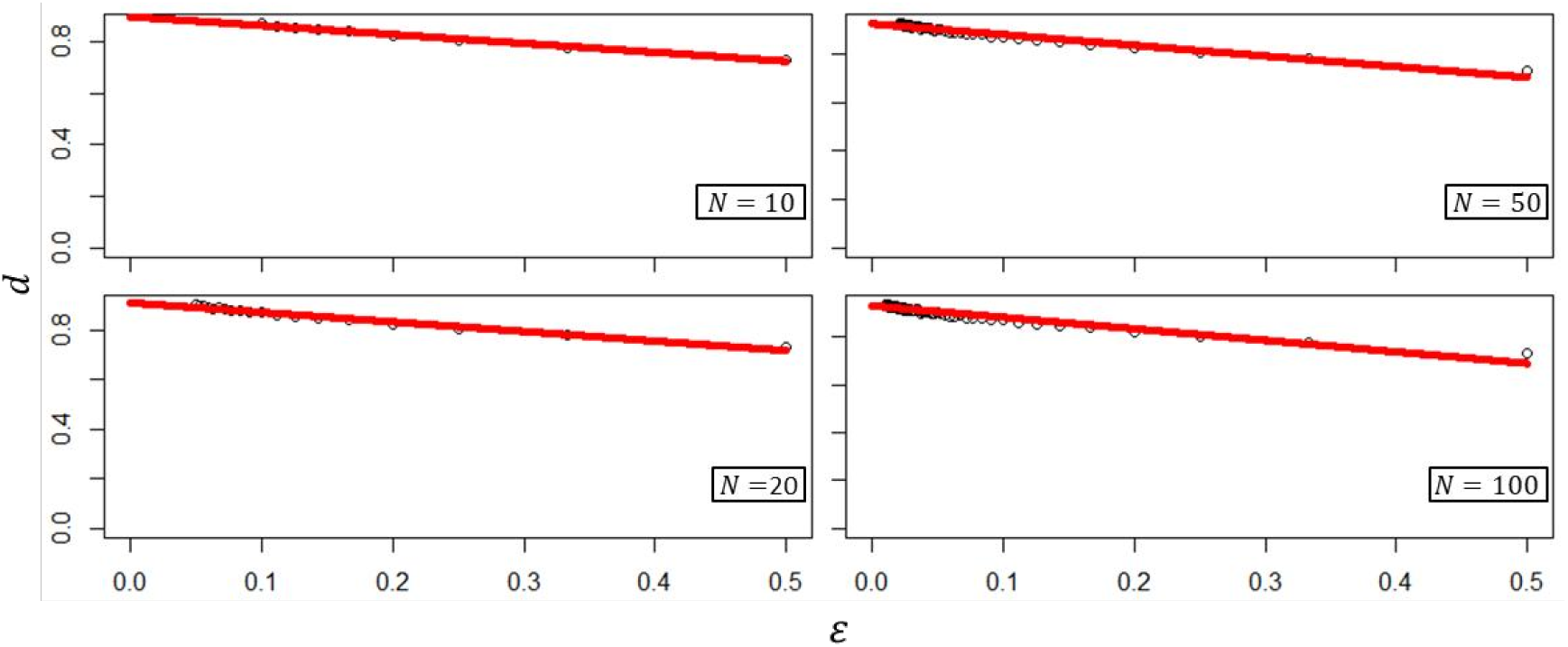
Shannon information dimension *d*_*ε*_ as function of the scaling parameter *ε* = 1*/N* for *N*_max_ = 10, 20, 50, 100. The input data is the same used in Fig.5 but the sites were sorted in increasing order with respect to *p*_comp_. Here the regression line (red line) no longer is horizontal.

**Figure 7:**
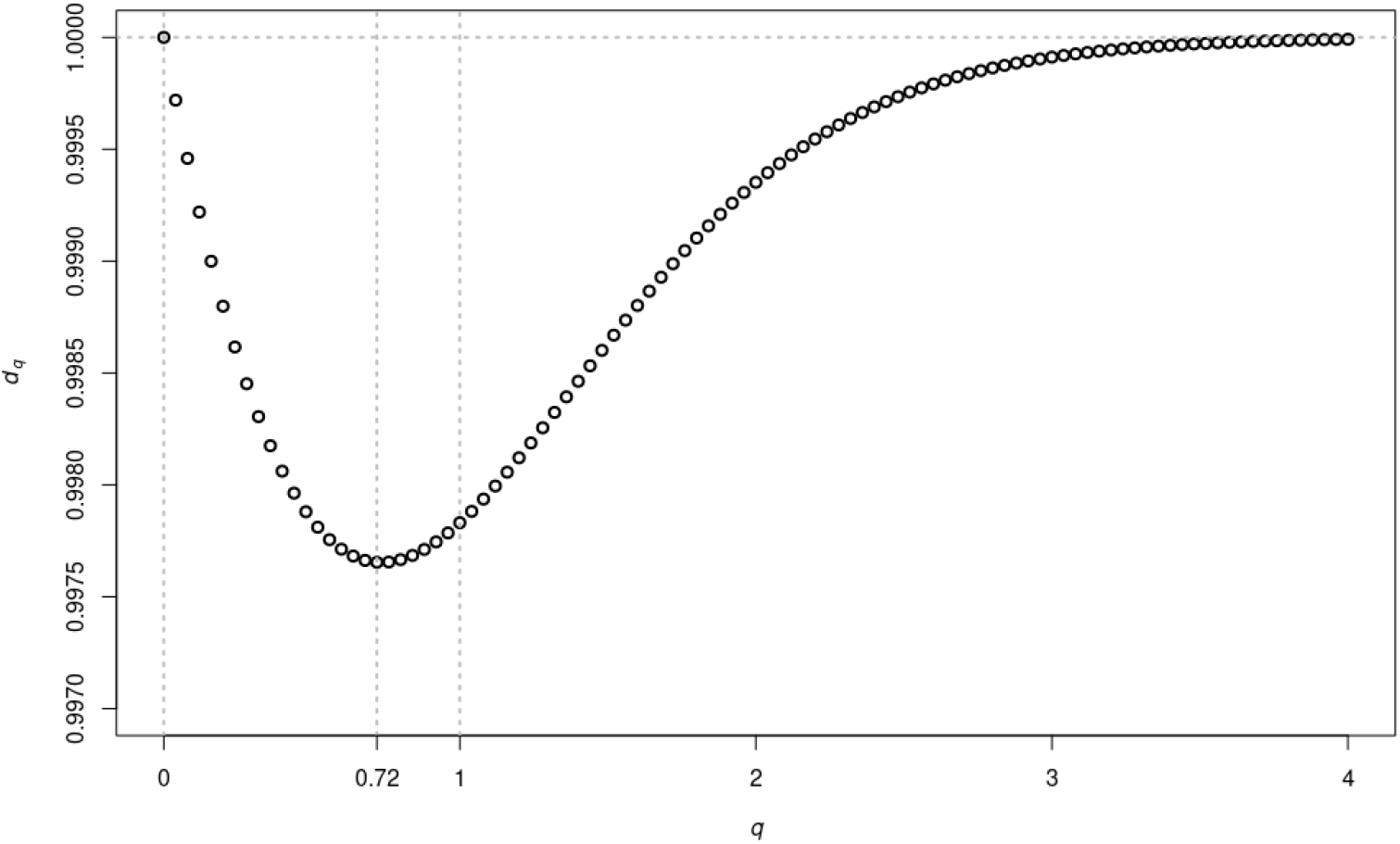
The *q*-logarithmic information dimension profile obtained from the same data as in Fig. 5 with *N*_max_ = 20. For *q* = 1 one obtains the Shannon information dimension *d* = *d*_1_ = 0.9978 as shown in Fig. 5. The value of *q* that makes *d*_*q*_ minimal is *q*^*^ = 0.72 with *d*_*q*_^*^ = 0.99765.

The second test consists of extending the notion of Shannon information dimension to the notion of *q*-logarithmic information dimension. This gives a whole family *d*_*q*_ of information dimensions, parametrized by a real parameter *q*, such that when *q* = 1 we recover the Shannon information dimension: *d*_1_ = *d*.

The advantage of defining a family of dimensions *d*_*q*_ is that rather than picking one *q* to work with, it’s best to consider all of them at once. The value of the parameter *q* reflects the strength of interactions between its components. When *q*≠ 1 the associated entropy is influenced by correlations or interactions between parts of the system. Thus, for a given system we construct *d*_*q*_ in a similar fashion as we did for the Shannon information dimension (see Supplementary Material, appendix **??** for details) and graph *d*_*q*_ against *q*. This is called a *dimension profile* of the system and is quite informative. Different values of the parameter *q* tell us different things about the system. Here, we consider only *q* ≥ 0 and note that there some values special values of *q* for which *d*_*q*_ is always the same: *d*_0_ = 1 and *d*_+∞_ = 1 (see Supplementary Material, appendix **??** for the definition of *d*_+∞_). Because of these properties, it follows that 0 *d*_*q*_ 1 for all *q* ≥ 0, i.e. *d*_*q*_ is bounded between 0 and 1. Furthermore, it follows that there is a *q*^*^ *>* 0 such that *d*_*q*_^*^ is minimal. The pair (*q*^*^, *d*_*q*_^*^) gives a ‘natural context’ for measuring the information of the system, where the correlations between the parts are maximally represented.

In Figure7 we show the *q*-logarithmic information dimension profile obtained from the same data as in Fig. 5 with *N*_max_ = 20. The fact that the Shannon information dimension *d* = *d*_1_ is continuously interpolated by *d*_*q*_ shows that, even though it is very close to 1, it is strictly smaller than 1. Moreover, the minimal dimension *d*_*q*_^*^ is attained when *q*^*^ = 0.72 showing that *q* = 1 is not the most natural *q*-index for the DMR.

## 4 Discussion

In this paper, we introduced the notion of the distribution of mutation rates (DMR) and showed that it can be estimated from sequencing data generated by an appropriate experimental design having a single replication round. The single replication round design allowed us to capture all mutations, including beneficial, neutral, deleterious, and/or lethal and yielded a substantial reduction of the bias from the effect of time on environmental selective pressure.

As mentioned in the introduction, there is a plethora of mutational rate values for several viruses, including HIV-1, HCV, and influenza virus, in the literature. Usually, these estimates were obtained by distinct methodologies providing a single numerical value a summarized mutational rate value, usually given a mean or a median.

The approach proposed here tries to leverage the wide coverage and depth furnished by high throughput sequencing technologies to estimate the DMR from sequencing data. Using the data generated by [33] we were able to successfully implement this approach by computing an empirical DMR for all 7 experimental conditions and the control considered by [33]. We also introduced a ‘consensus condition’ given by averaging over the 7 experimental conditions. Moreover, we were able to characterize the DMRs as being very good approximations of theoretical log-normal distributions.

Granted that the empirical DMRs were well-characterized distributions we proceeded further to study some of their basic properties. We computed some basic summary statistics of the empirical DMRs for all conditions and found a remarkable convergence of all central tendency measures: Mode, Median, and Mean (see Table 1).

Even though our focus here is to develop an approach centered on the DMR it is clear that it can be used to produce point and interval estimates of the mutation rates, for instance, using any one of the central tendency measures (Mode, Median, and Mean). Since the Mode and the Median are the most robust measures of central tendency – the Mean is very sensitive to variation in the tail of the distribution – the interval [Mode, Median] is a robust interval estimate for the mutation rate. Another possibility is to use the interval associated with the interquartile range (IQR), defined as [Q_1_, Q_3_], as a robust interval estimate for the mutation rate, given the large variability of the point-estimates of the mutation rate from the literature.

For example, from Table 1 we get that the IQR interval of the consensus condition is [3.7 × 10^−4^, 1.14 × 1 0^−3^] m/b/r. Now, this interval contains only a few of the several point estimates from the literature that were mentioned in the introduction. However, it is important to emphasize that such direct comparisons should be taken very carefully. First, most point estimates from the literature do not come with uncertainty quantification (e.g. standard error or confidence interval). Second, there are two units used to measure mutation rates: (i) mutations per base per cell infection (m/b/c) and (ii) mutations per base per replication cycle (m/b/r). They are not the same, since in a cell infection several replication cycles may occur, that is, a number of copying cycles per infected cell *r*_*c*_, which is a positive integer, satisfies *µ*_(*m/b/c*)_ */ µ*_(*m/b/r*)_ = *r*_*c*_. For HIV-1 we have that *r*_*c*_ = 1, since the template RNA is destroyed after the retro-transcription, but for other viruses, it is possible to have *r*_*c*_ *>* 1 [42]. Finally, we should mention that this apparent discrepancy is not surprising since it is expected that point/interval estimates obtained from the DMR tend to give higher values.

Despite the utility of the DMR to make point/interval estimates, it has much more potential to reflect the dynamics of a viral population than a single numerical value. By analyzing how the DMR behaves under down-scaling from the genome level to gene level and then to gene sub-fragments we identified a possible scale invariance of the DMR. The first consequence of this observation is that mutations spread all over the genome, not displaying any preferential hotspot. As seen from the summary statistics (Table 4), there is a remarkable convergence, indicating the almost complete position randomness of mutation occurrence.

One could argue that the *env* gene should display more mutations than other regions, considering that the envelope proteins are targets of neutralizing antibodies [25, 26]. Indeed, if the experiment design considered more than one replication cycle, the viral population could be exposed to some selection mechanism, which could be biased towards regions with more susceptibility. However, with a single replication cycle, any possibility of selection was shut down.

The emergence of the scale invariance suggests two questions: (i) is it possible to associate the presence of the apparent scale invariance of the DMR with some notion of information theory? (ii) can we show that the scale invariance has a fractal character?

Nature itself displays several fractal-like examples, such as protein complexes and the DNA [3, 28, 31, 44, 53]. The DNA could fit in the definition of ‘morphological fractal’ or ‘informational fractal’. A morphological fractal is defined by its geometry in terms of metric concepts, such as the Hausdorff dimension [21, 28]. The abstraction of the notion of fractal allows one to consider that the DNA exhibits a certain kind of selfsimilarity. By furthering the abstraction of the concept of fractal as ‘something’ that can be attributed to a notion of ‘dimension’ we are led to consider informational fractals. Moreover, it should be clear that any notion of ‘information dimension’ must be somehow associated with Shannon entropy. In fact, we have shown that the construction of a DMR introduced here, coupled with a down-scaling procedure can be used to define a notion of information dimension (based on Shannon entropy) to the system consisting of a genome together with its empirical DMR. As shown in Fig. 5, the information dimension is close to 1, independently of the fragment length. However, when the fragment length is close to 200 nucleotides (or below) the summary statistics start to show some divergence from the initial trend (Table 4).

An interesting question related to this minimal length where the downscaling breaks: is there a biological reason why the lower bound is around 200 nucleotides? Viral genomes are the most compressed and highly optimized self-replicating genomes, presenting few or no redundancy at all [22, 42]. Of course, such extreme genome reduction is one of the intracellular obligatory entities strategies to accomplish the self-replication process. For example, the smaller known viruses belong to the *Circoviridae* family, presenting about 1.7 to 2.2 thousand nucleotides. [4, 29]. The smallest unicellular organism is the *Mycoplasma genitalium* with a genome about 580 kb [15, 49]. In 2016, [19] introduced the JCVI-syn3.0, a self-replicating cell with the smallest synthetic bacterial genome of 531 kbp. However, viroids can display genomes of 200 to 400 nucleotides [23] and about 80% of the human exons are about 200 nucleotides [40]. These last two cases suggest that the minimal length that a genomic fragment carrying information is close to the lower bound of ~200 was found in our analysis.

The unusual robustness of the DMR shown in all our analyses leads us to a remarkable conclusion: the DMR constructed from the genome sequencing of HIV-1 is a ‘quasi-universal’ feature of this virus. In fact, as we mentioned before, the summary statistics of all 7 conditions, the control, and the consensus display sharp convergence in all statistics. The skewness estimation and the pairwise comparison using the two sample Kolmogorov-Smirnov test concurrently split the set of 7 conditions and the control into three groups: (i) the control, (ii) the stimulated conditions, and (iii) the non-stimulated conditions. Therefore, apart from the subtle difference between stimulated and non-stimulated conditions, the empirical DMRs obtained from the 7 conditions appeared to be essentially indistinguishable.

On a more speculative note, we would like to entertain possible explanations for the quasi-universality of the DMR. Recall that our experimental and analytical setup is designed to capture the rates of spontaneous mutations, which mainly occur due to errors made by DNA polymerases during replication. In a single-round infection event, with minimal selection pressure, the polymerase effects should be highlighted. Accordingly, any parameter associated with spontaneous mutations should be an intrinsic property of the replication machinery of the virus, independently of any exogenous influence.

When a DNA polymerase adds to the end (terminus) of a growing strand, sometimes a non-complementary unit can be added: this is often called a ‘misinsertion’ error. The resulting ‘terminal mispair’ can be corrected by a proofreading system if it exists, otherwise, it becomes an ‘internal mispair’. For example, in the *E. coli* polymerase III, misinsertions occur at a rate of 10^−4^ or 10^−5^ m/b/r for transition or transversion mispairs, respectively. The effectiveness of proofreading in this case is such that the rate of formation of internal mispairs is 50-fold to 100-fold lower than for terminal mispairs [27].

Nevertheless, most RNA viruses do not have any proofreading system (exceptions to this come from the order of Nidovirales, such as the Coronaviruses). Hence, the expected mutation rate should be in the range [10^−4^, 10^−5^] m/b/r. Since all point and interval estimates of mutation rates derived from the empirical DMR fall into this range, it is plausible to suppose that the empirical DMR captured the spontaneous mutation rates.

How does a misinsertion error originate? There are several non-exclusive possibilities. Errors may reflect ‘tautomeric shifts’ to alternative forms that favor non-canonical pairings: if the polymerase inserts a nucleotide while that nucleotide, or the template nucleotide, is in such an altered state, a non-canonical pairing is favored. It has been determined that tautomerism occurs at a frequency ~10^−4^ [51, 52, 54]. This is close to the spontaneous mutation rate reported here, but there is a problem: a tautomeric shift has to occur exactly at the same nucleotide that the polymerase is about to read. In any case, the joint probability of the simultaneous occurrence of these two events^1^ gives a (positive) lower bound to the spontaneous mutation rate, but it cannot exhaust the possibilities. A second possible mechanism is related to non-canonical ‘wobble’ pairings of canonical bases, due to the flexibility of the DNA helix, which gives it the capability to accommodate slightly misshaped pairings [27]. Finally, a third possible mechanism of misinsertion could be caused by conformational shifts due to the quantum tunneling of electrons. In particular, this means that ordinary replication errors may be subject to quantum indeterminacy [46]. It is very likely that in reality, these three mechanisms are at work simultaneously and non-independently (even in conjunction with others not mentioned above) to produce the spontaneous mutation rates that we observe.

## 5 Conclusions

In this paper, we have shown that it is possible to empirically estimate the distribution of mutation rates (DMR) from sequencing data. Furthermore, we were able the theoretically characterize the DMR as a log-normal distribution. By exploring the basic properties of the DMR we have shown that it can give point and/or interval estimates of mutation rates, that are reasonably close to estimates reported in the literature.

We have mentioned that the notion of DMR has been used in other papers [2, 50]. However, to the best of our knowledge, we seem to be the first to propose an estimation method for DMR from sequencing data. For that reason, we could not find any other experiment or sequencing dataset in the literature to which we could apply the methods of this paper.

The method proposed in this paper is based on two main ingredients: (i) a cell culture experimental design that provides exactly one replication cycle of a viral population and (ii) a method for estimation of mutation probabilities from high throughput sequencing data. We building on previous results on HIV-1 done by our group, where the item (i) above was accomplished in [33] and item (ii) in [56]. Nevertheless, the method can be in principle, applied to any virus, as long as some conditions are fulfilled: (1) as mentioned before, the viral population must undergo a single replication cycle. In the situation where it is not possible to ensure a single replication cycle, but only single cell infections, one should employ the appropriate unit of measurement. (2) one must have, samples of the genome before (at least one) and after (as many as desired) the replication cycle, and all samples must be sequenced. For instance, in our case, the sample obtained before the replication cycle was called, the control condition. (3) for each site in the assembled genome one must have an estimate of the probability of all four bases (p_A, p_T, p_G, and p_C, with p_A+p_T+p_G+p_C= 1). Granted these three conditions the construction of the empirical DMR follows as described before.

## Acknowledgements

The research of TNF was supported by Conselho Nacional de Desenvolvimento Científico e Tecnolǵico (CNPq) grant 140399/2020-8 and Fundação de Amparo à Pesquisa do Estado de São Paulo (FAPESP) grant 2024/08272-4. The research of IMVGC was supported by Fundação de Amparo à Pesquisa do Estado de São Paulo (FAPESP) grant 2021/11946-9.

## Research Ethics

This work was approved by the Research Ethics Committee of the Universidade Federal de São Paulo, registration #4.987.375 and certificate #50279321.7.0000.5505.

## Footnotes

1 Assuming that the genome has a length ∼10^4^ nucleotides and that both events are independent, the probability of simultaneous occurrence of both is ∼10^*−*8^ m/b/r.

