## Supplementary Material for "Enlarging viral mutation estimation: a view from the distribution of mutation rates"

March 20, 2025

#### Appendix: Supplementary Material

#### A Entropy and Information Dimension

The *Shannon entropy* of a finite probability distribution $P = \{p_i : i = 1, \ldots, k\}$, with $0 < p_i < 1$ and $\sum_{i=0}^{k} p_i = 1$, is given by [5]:

$$S = -\sum_{i=1}^{k} p_i \log p_i \tag{A.1}$$

In Shannon's original information-theoretic formulation, the logarithm was given in base 2 ($\log_2$), so that the Shannon entropy represents the expected number of yes/no questions necessary to determine an object's classification. Since then, it has become more common to use either the natural (base $e$) logarithm (log or ln) or the base 10 logarithm ($\log_{10}$). In this paper, we will always use the natural logarithm, denoted by 'log' and its corresponding exponential function denoted by 'exp'.

A coarse-grained approximation of a system is given by a partition of its state space $X$. A *partition* $\mathcal{A} = \{A_j : j = 1, \ldots, N\}$ of a set $X$ is a collection of nonempty, non-intersecting measurable sets $A_j$ that cover $X$. Thus, there is an element of the partition corresponding to every possible outcome of a measurement of the system [1]. Assume that a probability distribution $P_{\mathcal{A}} = \{p_j = p(A_j) : j = 1, \ldots, N\}$ is attached to the partition $\mathcal{A}$ – here $p$ is a function that associates a real number in $(0, 1)$ to each set $A_j$, such that $\sum_{j=0}^{N} p(A_j) = 1$. Then we define the entropy of the partition $\mathcal{A}$ by

$$S_{\mathcal{A}} = -\sum_{j=1}^{N} p_j \log p_j \tag{A.2}$$

Let $\mathcal{A}_N$ and $\mathcal{A}_M$ be finite partitions of the state space with $N < M$. We say that $\mathcal{A}_M$ is a *refinement* of $\mathcal{A}_N$ and write $\mathcal{A}_N \leq \mathcal{A}_M$ if every set in $\mathcal{A}_N$ is a union of sets in $\mathcal{A}_M$. Given sequence of partitions

[1]Lab. Virologia, Instituto Butantan, São Paulo, SP, Brazil.
[2]Departamento de Microbiologia, Imunologia e Parasitologia, Departamento de Medicina, Laboratório de Retrovirologia, Universidade Federal de São Paulo, São Paulo, SP, Brazil.
[3]Centro de Bioinformática Médica, Universidade Federal de São Paulo, São Paulo, SP, Brazil.
[†]These authors share the first authorship.
[‡]These authors share the senior authorship.
[*]Corresponding author.

$\mathcal{A}_1 \leq \mathcal{A}_2 \leq \ldots \leq \mathcal{A}_N$ of the state space $X$, one can consider a sequence of probability distributions $P_{\mathcal{A}_N}$ and the corresponding sequence of entropies $S_{\mathcal{A}_N}$. The *information dimension* of the system is defined by

$$d = \lim_{N\to+\infty} \frac{-\sum_{j=1}^{N} p_j \log p_j}{\log N} = \lim_{N\to+\infty} \frac{S_{\mathcal{A}_N}}{\log N} \tag{A.3}$$

Note that the distributions $P_{\mathcal{A}_N}$ are univariate, hence it can be shown that $0 \leq d \leq 1$ [4].

The intuition behind this definition is the following. Suppose that we have a notion of "size" of the sets $A_j$ of a partition $\mathcal{A}_N$ and that each set has approximately the same "size", $\varepsilon \approx 1/N$. In this setting, each set $A_j$ is called a *box*, and thus $\varepsilon$ is the "box size" parameter. Then we can write

$$d \approx \frac{S(\varepsilon)}{-\log \varepsilon} \qquad \text{for} \quad \varepsilon \to 0^+ \tag{A.4}$$

where $S(\varepsilon) = S_{\mathcal{A}_N}$ is the average information per box. Hence, the information dimension $d$ is the exponent of the box size scaling of the average information needed to identify an occupied box, that is

$$\exp S(\varepsilon) \approx \varepsilon^{-d} \qquad \text{for} \quad \varepsilon \to 0^+ \tag{A.5}$$

The exponential of the Shannon entropy $\exp S$ appears in many fields and thus has different names. It is called *perplexity* in the fields of machine learning and statistical modeling and is a measure of uncertainty in the value of a sample from a discrete probability distribution [2]. The larger the perplexity, the less likely it is that an observer can guess the value that will be drawn from the distribution. It is called *diversity index of order* 1 in the fields of ecology and demography [3]. It is a measure of how many different types (e.g., species) there are in a dataset, i.e., the "effective number of species" in a community.

Shannon entropy is fundamental, but it is not the only useful or natural notion of entropy, even in the context of a single probability distribution on a finite set. Two one-parameter families of entropies include Shannon entropy as a member: the Rényi entropies and the $q$-logarithmic or Tsallis entropies [3, Figure 4.1]. Both are indexed by a real parameter $q$, and both have Shannon entropy as the case $q = 1$. Varying $q$ away from 1 can be thought of as deforming Shannon entropy. As in other mathematical contexts where the word 'deformation' is used, the undeformed object (Shannon entropy) has uniquely good properties that are lost after deformation, but the deformed objects nevertheless retain some of the original object's features.

Historically, the first deformation of Shannon entropy was the Rényi entropy family, introduced by Rényi in 1961. The exponential of the Rényi $q$-entropy is known in ecology as the 'Hill number of order $q$', denoted by $D_q$, introduced by Mark Hill in 1973 (acknowledging the prior work of Rényi). In particular, $D_1 = expS$ is the exponential of the Shannon entropy. There is a precise mathematical sense in which the Hill numbers are the only well-behaved measures of diversity [3].

The $q$-logarithmic entropies have been discovered and rediscovered repeatedly. They seem to have first appeared in a 1967 paper on information and classification by Havrda and Charvát. They were rediscovered in a 1988 paper on statistical physics by Tsallis (without reference to any of the previous rediscoverers). It was after Tsallis that they gained widespread recognition and became most commonly named. The term '$q$-logarithmic entropy' is new but has the benefits of being descriptive and of not perpetuating a misattribution [3, Remark 4.1.4].

The $q$-logarithmic and Rényi entropies have exactly the same content: for each finite value of $q$, there is a simple formula for one of them in terms of the other, and vice versa. But they have different and complementary algebraic properties. While the Rényi entropies are used to define the diversity indices of order $q$, the $q$-logarithmic entropies are used to define the information dimension of order $q$.

Let be given a finite probability distribution $P = \{p_i : i = 1, \ldots, k\}$, with $0 < p_i < 1$ and $\sum_{i=0}^{k} p_i = 1$ and let $q \in \mathbb{R}$ be a parameter. The *$q$-logarithmic entropy* of $P$ is defined as

$$S_q = \begin{cases} -\sum_{i=1}^{k} p_i \log p_i & \text{if} \quad q = 1 \\ \frac{1}{q-1}(1 - \sum_{i=1}^{k} p_i^q) & \text{if} \quad q \neq 1 \end{cases} \tag{A.6}$$

and the *Rényi entropy* of $P$ is defined as

$$H_q = \begin{cases} -\sum_{i=1}^{k} p_i \log p_i & \text{if} \quad q = 1 \\ \frac{1}{1-q} \log(\sum_{i=1}^{k} p_i^q) & \text{if} \quad q \neq 1 \end{cases} \tag{A.7}$$

Note that both $S_q$ and $H_q$ coincide with Shannon entropy when $q = 1$: $H_1 = S_1 = S$. Moreover, they are both continuous with respect to $q$, as it can be shown that $\lim_{q \to 1} H_q = \lim_{q \to 1} S_q = S$ (see [3]). Furthermore, the limits $q \to \pm\infty$ always exist and this allows one to define $S_q$ and $H_q$ for $q = \pm\infty$ (see [3]). The relation between $S_q$ and $H_q$ is

$$\exp(H_q) = \exp_q(S_q) \tag{A.8}$$

where the function $\exp_q$ is the *q-deformed exponential*, defined as

$$\exp_q(u) = \begin{cases} \exp(u) & \text{if} \quad q = 1 \\ (1 + (1-q)u)^{\frac{1}{1-q}} & \text{if} \quad q \neq 1 \end{cases} \tag{A.9}$$

In principle, the parameter $q$ can be any real number, but from now on we consider only the case $q \geq 0$.

Let be given a system $X$ equipped with coarse-grained approximation on its state space defined by a sequence of partitions $\mathcal{A}_1 \leq \mathcal{A}_2 \leq \ldots \leq \mathcal{A}_N$. Then we can proceed as before and construct a sequence of probability distributions $P_{\mathcal{A}_N,q}$ and the corresponding sequence of entropies $S_{\mathcal{A}_N,q}$ (defined as in A.2, using formula A.6 instead of A.1). The *information dimension* of order $q$ ($q \geq 0$) of the system $X$ is defined by

$$d_q = \lim_{N \to +\infty} \frac{S_{\mathcal{A}_N,q}}{\ln_q N} \tag{A.10}$$

Her, the function $\ln_q$ is the *q-deformed logarithm*, defined as

$$\ln_q(u) = \begin{cases} \ln(u) & \text{if} \quad q = 1 \\ \dfrac{u^{(1-q)} - 1}{1 - q} & \text{if} \quad q \neq 1 \end{cases} \tag{A.11}$$

Again, since the distributions $P_{\mathcal{A}_N,q}$ are univariate, it follows that $0 \leq d_q \leq 1$, for $q \geq 0$.

Since $q \geq 0$, the limit cases are given by $q = 0$ and $q = +\infty$ that are as follows

$$d_0 = 1 \qquad \text{and} \qquad d_{+\infty} = \lim_{q \to +\infty} d_q = 1 \tag{A.12}$$

Now, using relation (A.8) and the fact that the exponential of the Rényi (Shannon) entropy $\exp H_q$ ($\exp S$) is the diversity index or Hill number of order $q$ (1), denoted by $D_q$, one can show the following deformed version of (A.5): the information dimension $d_q$ is the exponent of the *q-deformed box size scaling* of the average *Rényi* information needed to identify an occupied box, that is

$$\exp H_q(\varepsilon) = \exp_q S_q(\varepsilon) \approx [\varepsilon^{-d_q}]_q \qquad \text{for} \quad \varepsilon \to 0^+ \tag{A.13}$$

Again, here $S_q(\varepsilon) = S_{\mathcal{A}_N,q}$ and $H_q(\varepsilon) = H_{\mathcal{A}_N,q}$ is the average Rényi information per box, defined as before. The *q-deformed scaling* is defined by $[\varepsilon^{-d_q}]_q = \exp_q(-d_q \ln_{(2-q)}(\varepsilon))$ (see [3]).

### B Supplementary Figures

In this appendix, we present three sets of plots. The first (Figure 1) shows the Histograms of the log-transformed empirical distributions of mutational rates for the 7 seven experimental conditions and the control. The second (Figure 2) shows the Q-Q plots of the log-transformed empirical distributions of mutational rates for the 7 seven experimental conditions and the control. The third (Figure 3) shows the gaps after excluding sites for the 7 seven experimental conditions and the control.

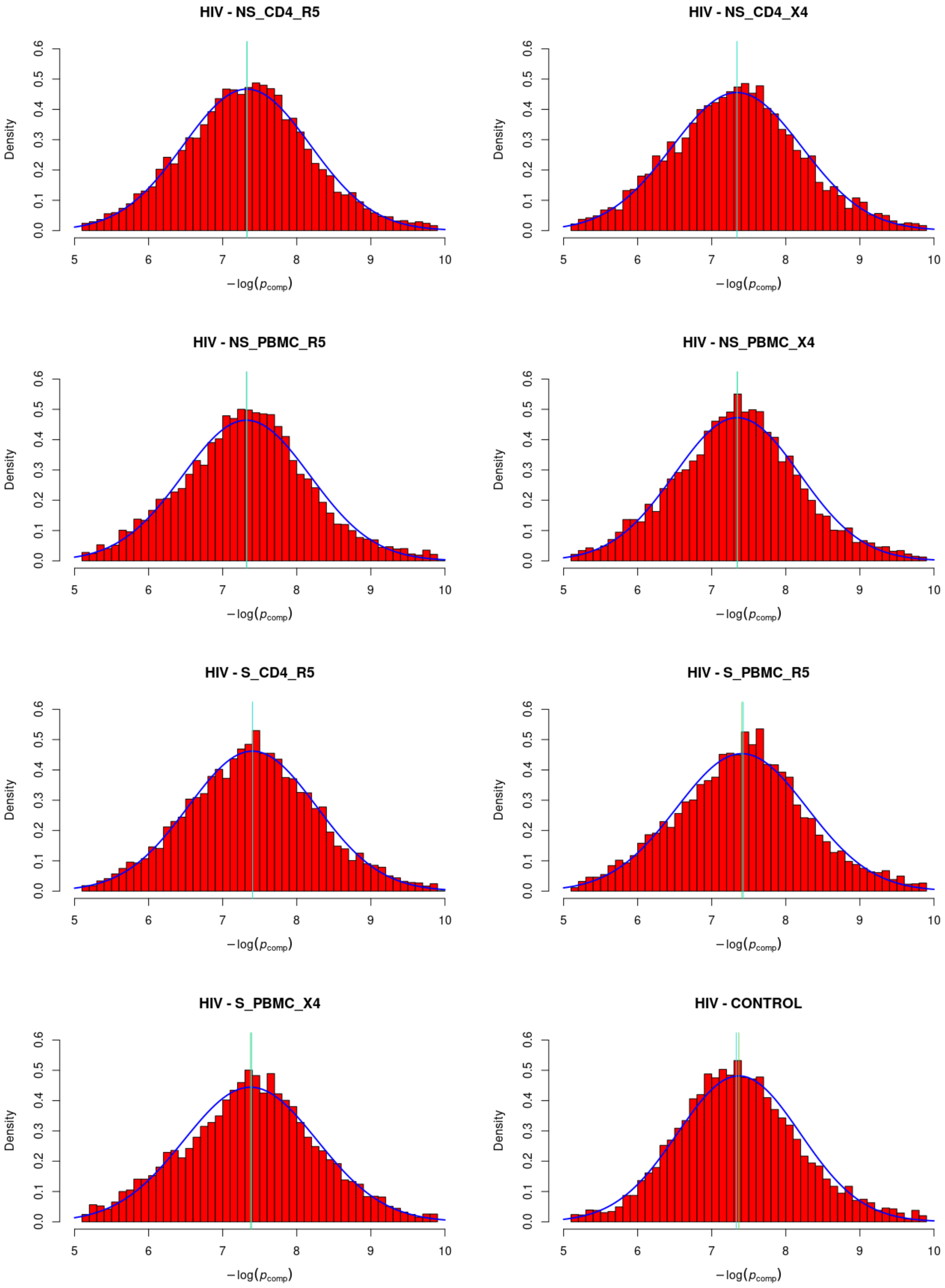


Figure 1: Histograms of the log-transformed empirical distributions of mutational rates for the 7 seven experimental conditions and the control.

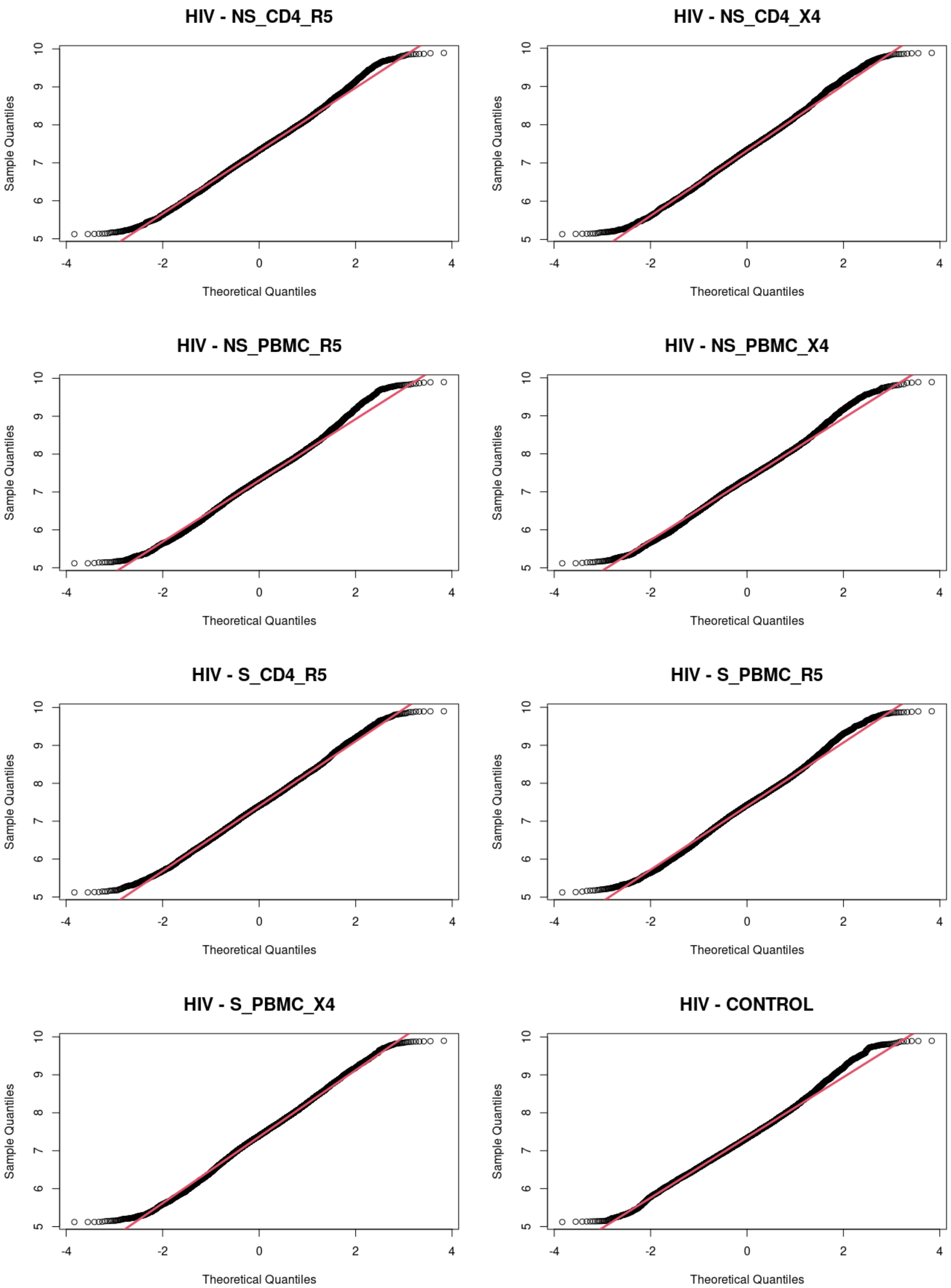


Figure 2: Q-Q plots of the log-transformed empirical distributions of mutational rates for the 7 seven experimental conditions and the control.

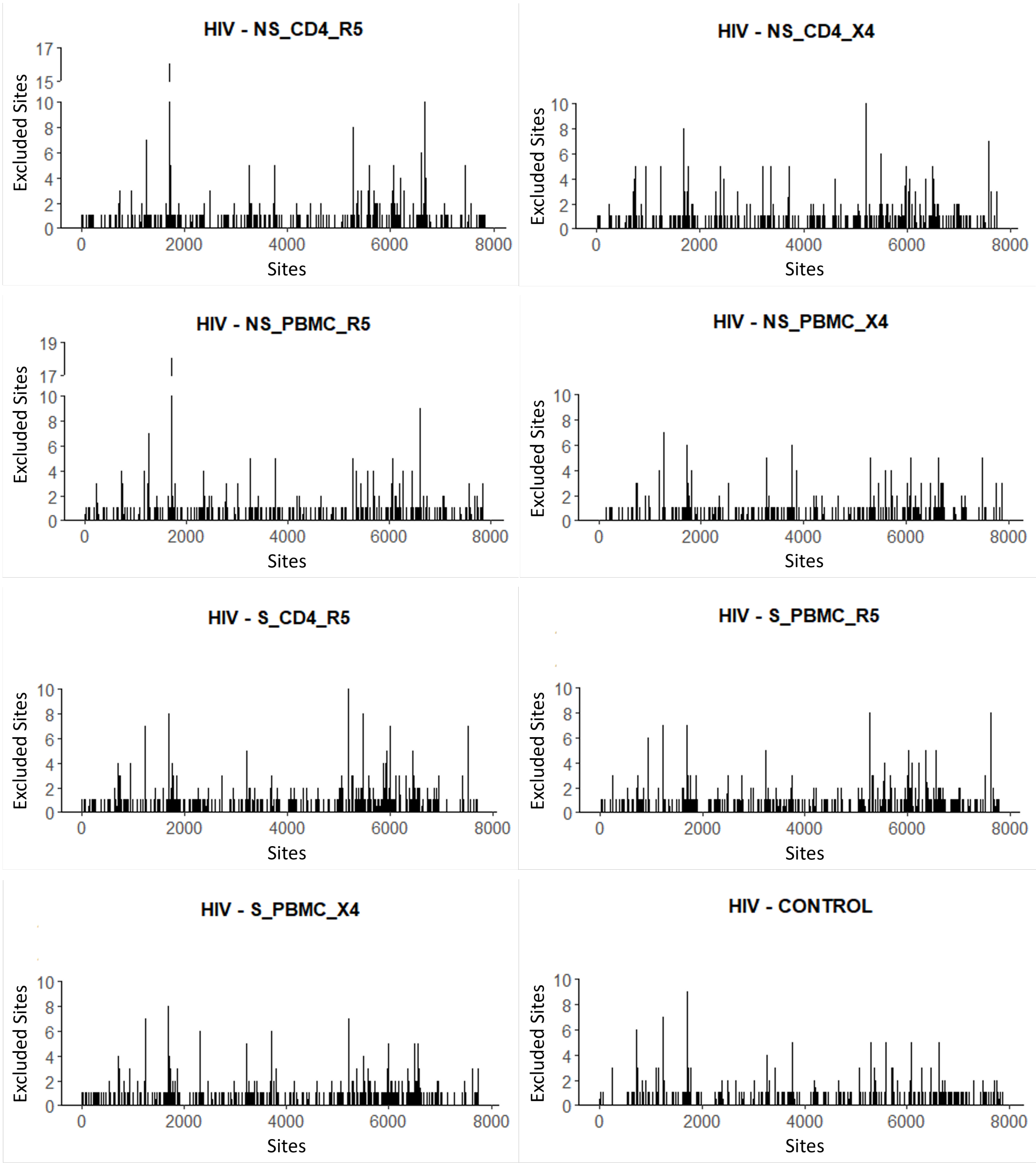


Figure 3: Gaps positions for the 7 seven experimental conditions and the control. Each bar is the size of the gap after each remaining site. The marks in the $x$-axis represent the absolute number of sites in the specific sequence and not their location in the genome.
